# Structural predictions of the SNX-RGS proteins suggest they belong to a new class of lipid transfer proteins

**DOI:** 10.1101/2021.11.30.470681

**Authors:** Blessy Paul, Saroja Weeratunga, Vikas A. Tillu, Hanaa Hariri, W. Mike Henne, Brett M. Collins

**Affiliations:** The University of Queensland, Institute for Molecular Bioscience, Queensland, 4072, Australia; Department of Cell Biology, University of Texas Southwestern Medical Center, Dallas, TX 75390, United States

**Keywords:** sorting nexin (SNX), regulator of G-protein signaling (RGS), lipid droplets (LDs), AlphaFold2, X-ray crystallography, lipid transfer protein (LTP), phox homology (PX) domain

## Abstract

Recent advances in protein structure prediction using machine learning such as AlphaFold2 and RosettaFold presage a revolution in structural biology. Genome-wide predictions of protein structures are providing unprecedented insights into their architecture and intradomain interactions, and applications have already progressed towards assessing protein complex formation. Here we present detailed analyses of the sorting nexin proteins that contain regulator of G-protein signalling domains (SNX-RGS proteins), providing a key example of the ability of AlphaFold2 to reveal novel structures with previously unsuspected biological functions. These large proteins are conserved in most eukaryotes and are known to associate with lipid droplets (LDs) and sites of LD-membrane contacts, with key roles in regulating lipid metabolism. They possess five domains, including an N-terminal transmembrane domain that anchors them to the endoplasmic reticulum, an RGS domain, a lipid interacting phox homology (PX) domain and two additional domains named the PXA and PXC domains of unknown structure and function. Here we report the crystal structure of the RGS domain of sorting nexin 25 (SNX25) and show that the AlphaFold2 prediction closely matches the experimental structure. Analysing the full-length SNX-RGS proteins across multiple homologues and species we find that the distant PXA and PXC domains in fact fold into a single unique structure that notably features a large and conserved hydrophobic pocket. The nature of this pocket strongly suggests a role in lipid or fatty acid binding, and we propose that these molecules represent a new class of conserved lipid transfer proteins.

## Introduction

The sorting nexins (SNXs) are a large family of proteins with diverse structures and functions. They are found in all eukaryotes and their defining feature is the presence of a phox homology (PX) domain, which most commonly associates with phosphatidylinositol phospholipids (PtdInsPs) to mediate interactions with membranes of the endolysosomal system ^1–3^. SNX proteins are typically grouped into sub-families based on the presence of additional functional domains such as membrane tubulating bin/amphiphysin/rvs (BAR) domains, protein interacting SH3 domains, and GTPase activating protein (GAP) domains among many others ^3^.

One of the most highly conserved sub-families are the SNX proteins with regulator of G-protein signalling (RGS) domains or SNX-RGS proteins ^4, 5^. In humans there are four genes encoding SNX-RGS proteins, SNX13, SNX14, SNX19 and SNX25. Other well-characterised family members include Mdm1 from *Saccharomyces cerevisiae* and Snazarus (Snz) from *Drosophila melanogaster*. These proteins have relatively low sequence similarity across homologues and across species but share a conserved architecture with an N-terminal hydrophobic membrane anchor, and a central RGS domain and PX domain (except for SNX19 orthologues which lack the RGS domain). In addition, the RGS and PX domains are flanked by two sequences referred to as the PX-associated domains PXA and PXC at the N- and C-terminus respectively (**Fig. 1A**). The N-terminal membrane anchor mediates localisation to the endoplasmic reticulum (ER) ^6–10^, and the proteins typically localise to sites of new lipid droplet synthesis and can enhance membrane tethering of the ER to other membrane compartments via their PX domains including the vacuole in yeast ^8, 11, 12^, the plasma membrane in *Drosophila* ^10^ and endolysosomal compartments in mammalian cells ^6, 7, 9, 13^.

**Figure 1.**
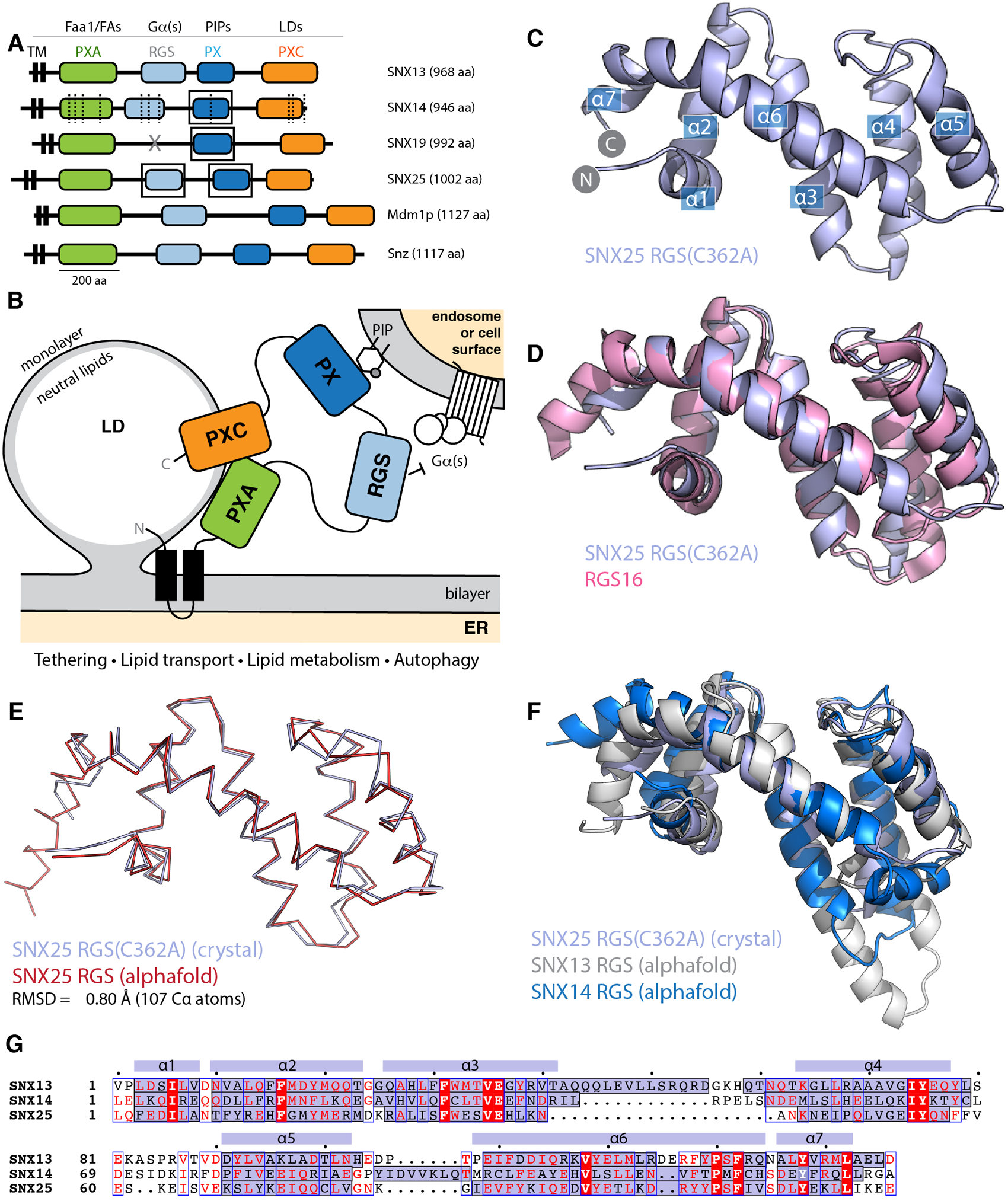
The SNX-RGS protein family. **(A)** Domains of human, *Drosophila*, and *S. cerevisiae* SNX-RGS family members. Putative ligands for each domain are indicated above. SNX14 mutations causing cerebellar ataxia are shown with dashed lines ^14, 15^. Boxes indicate domains with experimental structures determined. (**B**) The current working model for the architecture, localisation and function of the SNX-RGS protein family and their potential roles at the interface between the ER, LDs, and membranes of the endolysosomal system. (**C**) Crystal structure of human SNX25(C362A) RGS domain in cartoon diagram. Helical secondary structure elements are indicated. (**D**) The crystal structure of human SNX25(C362A) RGS domain (light blue) is aligned with the canonical RGS domain of RGS16 (pink)(PDB ID 2IK8)^28^ and shows the expected topology. (**E**) Cα ribbon diagram showing the overlay of the SNX25(C362A) RGS domain (light blue) with the same sequence predicted in the AlphaFold2 database (red) (Q9H3E2). (**F**) Cartoon representation of SNX25(C362A) RGS domain compared with the SNX13 and SNX14 RGS domains from the AlphaFold2 database (Q9Y5W8; Q9Y5W7). (**G**) Sequence alignment of human SNX13, SNX14 and SNX25 RGS domains based on structural comparisons. The secondary structure of SNX25 derived from its crystal structure is shown schematically above the alignment, while the α-helical regions of the three proteins from their AlphaFold2 predictions are shaded blue within the alignment.

The precise function of the SNX-RGS proteins is still unclear although their mutation or depletion has significant impacts on cellular physiology, particularly with respect to lipid homeostasis and endolysosomal function. Mutations in human SNX14 cause the autosomal-recessive cerebellar ataxia and intellectual disability syndrome SCAR20, with a common cellular phenotype of autophagic structures containing undigested material and increased cholesterol accumulation in endolysosomal organelles ^6, 14, 15^. In mice SNX13, SNX14 and SNX25 are all essential for normal development ^16–18^, while altering the expression of Snz significantly extends the lifespan of flies ^10, 19^. Previous work has defined the structures and lipid binding properties of the PX domains of these proteins; the human SNX13 and SNX19 PX domains bind the endosomal lipid PtdIns3P and SNX25 associates with multiple phosphorylated PtdInsP species, while SNX14 has little affinity for PtdInsP lipids due to an altered binding pocket ^1, 5, 10^. The RGS domains of SNX13 and SNX14 associate with the Gα_s_ subunit of trimeric G-proteins ^20, 21^. While SNX13 was originally discovered as a GTPase activating protein (GAP) for Gα_s_ ^21^, SNX14 can bind Gα_s_ but has so far been found to not stimulate its GAP activity^20^. SNX13 and SNX14 have also been found to associate with the endosomal Rab GTPase Rab9A, although the functional significance of this is unclear ^22^. Proteomics studies have identified association of yeast Mdm1 with Faa1 long-chain-fatty-acid--CoA ligase, while *Drosophila* Snz and human SNX14 were associated with fatty acid desaturases Desat1 and SCD1 respectively ^10, 23^, suggesting a role in lipid and fatty acid regulation. In line with this, members of the SNX-RGS protein family have almost universally been shown to localize to ER-lipid-droplet (LD) contact sites with other organelles (**Fig. 1B**). Yeast Mdm1 localizes to nucleus-LD-vacuole junctions^8, 12, 24^. Human SNX14 localizes to ER-LD contacts^7^, and SNX19 has recently been shown to localize to ER-lysosome contacts, where it also contacts LDs ^9^.

A better understanding of the function(s) of the SNX-RGS proteins requires further knowledge of their structures and molecular interactions. Here we have solved the crystal structure of the human SNX25 RGS domain, revealing a typical α-helical RGS fold. Building on this we took advantage of the recently developed AlphaFold2 machine learning (ML)-based structural prediction algorithm ^25^ and the related AlphaFold2 database ^26^ to examine the structures of the full-length SNX-RGS proteins. These analyses revealed an intriguing structural fold not previously seen, that is formed by intramolecular association of the distal PXA and PXC domains. A remarkable feature of this structure is the presence of a large hydrophobic cavity. Based on their structures and known localisation and interactions we propose that the SNX-RGS proteins may represent a previously undefined class of lipid transfer proteins (LTPs) that are likely to mediate binding of multiple lipids or other fatty acid derived molecules. Further, using AlphaFold2 we identify an additional yeast PX domain-containing protein Lec1/Ypr097w that contains another likely lipid-binding structure that shares superficial resemblance but is not directly related to the SNX-RGS proteins. This work provides a new understanding of the likely function of the SNX-RGS proteins, presents a structural bioinformatics comparison of novel lipid binding domains across various eukaryotic species, and further demonstrates the utility of new ML-based structure prediction programs such as AlphaFold2 to develop novel structural and functional insights.

## Results

### Crystal structure of the human SNX25 RGS domain

The SNX-RGS family members are typically represented as modular proteins with four to five distinct conserved domains (**Fig. 1A and 1B**) ^4, 5, 8, 9, 12, 14, 15, 19, 21, 23, 27^. Our previous studies examined the structures and phosphoinositide binding properties of the PX domains of SNX14, SNX19 and SNX25 using NMR and X-ray crystallography ^1, 5, 10^. Here we have also solved the structure of the human SNX25 RGS domain by X-ray crystallography in two crystal forms (**Fig. 1C**; **Fig. S1**; **Table S1**). The wildtype sequence of the RGS domain crystallises with a non-native disulfide bond formed by Cys526 between adjacent monomers in the crystal lattice which distorts the orientation of the helix α5. Mutation of Cys526 to alanine (C526A) prevents this bond formation and allows the domain to crystallise with a more typical α5 conformation similar for example to a canonical RGS domain such as RGS16 (PDB ID 2IK8)^28^ (**Fig. 1D**).

We next investigated how similar the experimentally determined SNX25 RGS domain was to the predicted structure from the AlphaFold2 database ^26^. The structure predicted by AlphaFold2 is remarkably close to the SNX25 crystal structure, with an overall root-mean-squared-deviation (RMSD) of 0.8 Å over 107 C_α_ atoms (**Fig. 1E**). Side chain conformations in general were likewise closely aligned between the experimental and predicted structures (**Fig. S2**). The predicted structures of the human SNX13 and SNX14 RGS domains from the AlphaFold2 database are generally similar to the experimental SNX25 RGS domain structure as expected, but with some notable differences (**Fig. 1F**). In general, the three domains have relatively low sequence homology, and this is reflected for example in a significantly longer α3-α4 extension in SNX13 compared to the other two proteins and (**Fig. 1F and 1G**). The experimental structures of the PX domains of SNX14, SNX19 and SNX25 were also compared with their respective AlphaFold2 predictions and in general, the AlphaFold2 models were found to be highly similar with only minor differences in sidechain positions and in some flexible loop regions. (**Fig. S2**). Overall, these comparisons confirm that the AlphaFold2 predictions of selected SNX-RGS domains are accurate models comparable to their known experimental structures, including the RGS domain of SNX25 for which no detailed structural information was available prior to this study.

### AlphaFold2 predictions of the yeast, fly and human SNX-RGS proteins reveal conserved PXA and PXC domain intramolecular interactions

Despite these structural insights into the SNX-RGS proteins, the structures of the N-terminal PXA and PXC domains bear no obvious sequence similarity to other proteins and their structures have not been determined, and the overall architecture of the proteins is unknown. We have so far been unsuccessful in expressing and purifying either full-length SNX-RGS proteins or any of their individual PXA and PXC domains in suitable quantities for structural or biochemical analyses. As the recently released AlphaFold2 database ^26^ includes predictions of the proteomes of multiple eukaryotic species from yeast to humans, we took advantage of this to examine the predicted structures of full-length SNX-RGS proteins across different species (**Fig. 2**; **Fig. S2**; **Fig. S3**).

**Figure 2.**
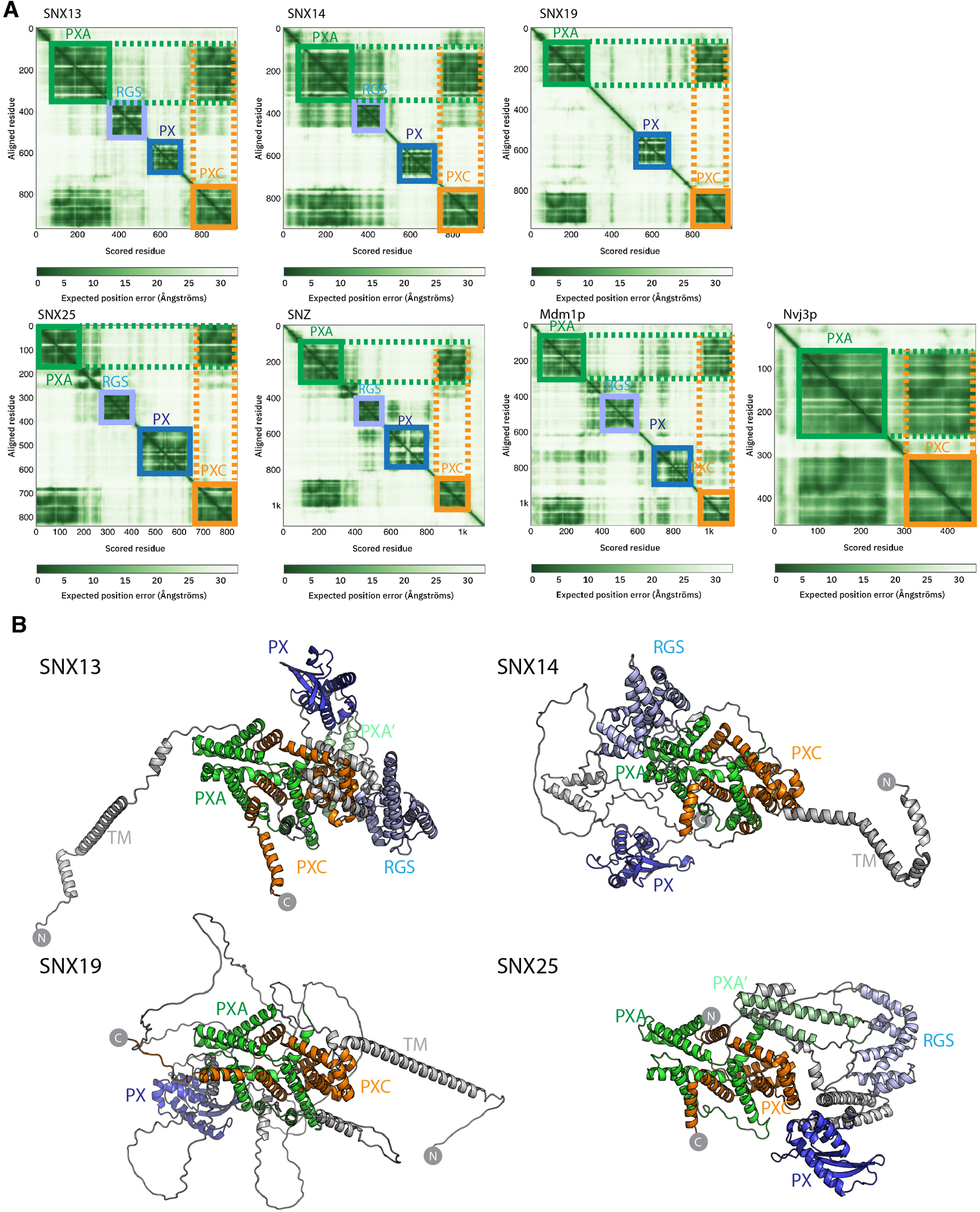
Structures of the SNX-RGS proteins predicted by AlphaFold2. **(A)** Predicted Alignment Error (PAE) plots from the AlphaFold2 database ^26^ are shown for human fly and yeast SNX-RGS proteins. In these plots all SNX-RGS proteins show a strong degree of correlation between the PXA and PXC domain suggesting these two domains are physically associated. (**B**) The predicted structures of human SNX-RGS proteins from the AlphaFold2 database. The PXA domain is coloured green, RGS domain in light blue, PX domain in blue and PXC domain in orange. The predicted TM domain and any unstructured linker regions are coloured grey. The structures are shown in the same orientation after alignments based on the PXA and PXC core region. This shows that the two domains are intimately entwined with each other, whereas the TM, RGS and PX domains are predicted to have flexible orientations relative to these domains.

The plots of the predicted alignment errors (PAE) of the AlphaFold2 models provide important insights into the organisation of the full-length proteins (**Fig. 2A**). The PAE measures AlphaFold’s expected position error at residue ‘x’ if the predicted and true structures were aligned on residue ‘y’. Square regions of the plot along the diagonal indicate the presence of globular domains where residues are structurally correlated with each other, while off-diagonal low PAE regions for residue pairs from two different domains is highly indicative of stable inter-domain interactions. The PAE plots for the human SNX-RGS proteins display expected low PAE regions for each of the four cytoplasmic domains downstream of the N-terminal ER membrane anchor (except for SNX19 which lacks an RGS domain). Notably, even though the N-terminal PXA domain and C-terminal PXC domains are around 500 amino-acids apart in the primary sequence, these two domains are invariably predicted to be tightly correlated with each other. Further, *Drosophila* Snz and yeast Mdm1p homologues show a similar high degree of structural correlation between their PXA and PXC domains.

The AlphaFold2 predicted atomic structures of the human SNX-RGS proteins are shown in **Fig. 2B** and **Fig. S3**. Consistent with secondary structure predictions ^5^ both the PXA and PXC domains are entirely α-helical, and as indicated by the PAE plots they are tightly associated with each other. Indeed, they form an overall structure with a highly entwined topology, which is structurally similar in all four proteins. Relative to this ‘core’ structure the N-terminal membrane anchor, and intervening RGS and PX domains adopt diverse orientations pointing to a substantial degree of structural dynamics in the full-length proteins. Note that the human SNX25 sequence in the AlphaFold2 database (Uniprot ID Q9H3E2) lacks the N-terminal membrane anchor as previously identified ^5, 29^ (Uniprot A0A494C0S0). We therefore also performed predictions of the full SNX25 sequence using the online ColabFold pipeline ^30^, which show a similar structure but with the additional N-terminal α-helical membrane anchor (**Fig. S4**). As for predictions across the family, the three separate models of human SNX25 generated by ColabFold suggest that the transmembrane anchor, RGS and PX domains adopt relatively flexible orientations with respect to the core PXA-PXC structure. Although not shown, both full-length Mdm1p and Snz have analogous architectures with a core structure composed of the PXA and PXC domains that adopt a flexible orientation with respect to the transmembrane, RGS and PX domains. Overall, the predicted structures of the RGS-SNX proteins are highly consistent across homologues and species and reveal a novel intramolecular association between the previously uncharacterised PXA and PXC domains.

### The PXA and PXC domains form an interwoven structure with a large conserved hydrophobic channel

For a closer examination of the PXA-PXC structure we focus on these domains from the SNX13 model (**Fig. 3A**; **Movie S1**). Both the PXA and PXC domains are entirely α-helical as predicted ^5^ and do not share any obvious structural similarity with previously characterised proteins. A notable feature of their structures is that they are intimately intertwined in such a way that neither domain would be expected to form their correct fold independently of the other. The threading of their α-helical structures together forms an overall elongated structure with a length of around 80 Å. The same α-helical topology and pattern of threading between the PXA and PXC domains is highly conserved and is seen in all SNX-RGS family members (**Fig. 3B**). Although the yeast protein Nvj3p (Ydr179w-a) is not found in higher eukaryotes, it was previously proposed to share weak sequence similarity with the PXA domain of the SNX-RGS proteins ^8^. Interestingly the AlphaFold2 prediction of Nvj3p shows a very similar structure to the combined PXA and PXC fold of the SNX-RGS proteins (**Fig, 3C**). Although it lacks a transmembrane, RGS or PX domains, and has only a minimal linker sequence between the two halves of its structure, it shares the same overall topology and is a bona fide PXA-PXC structure.

**Figure 3.**
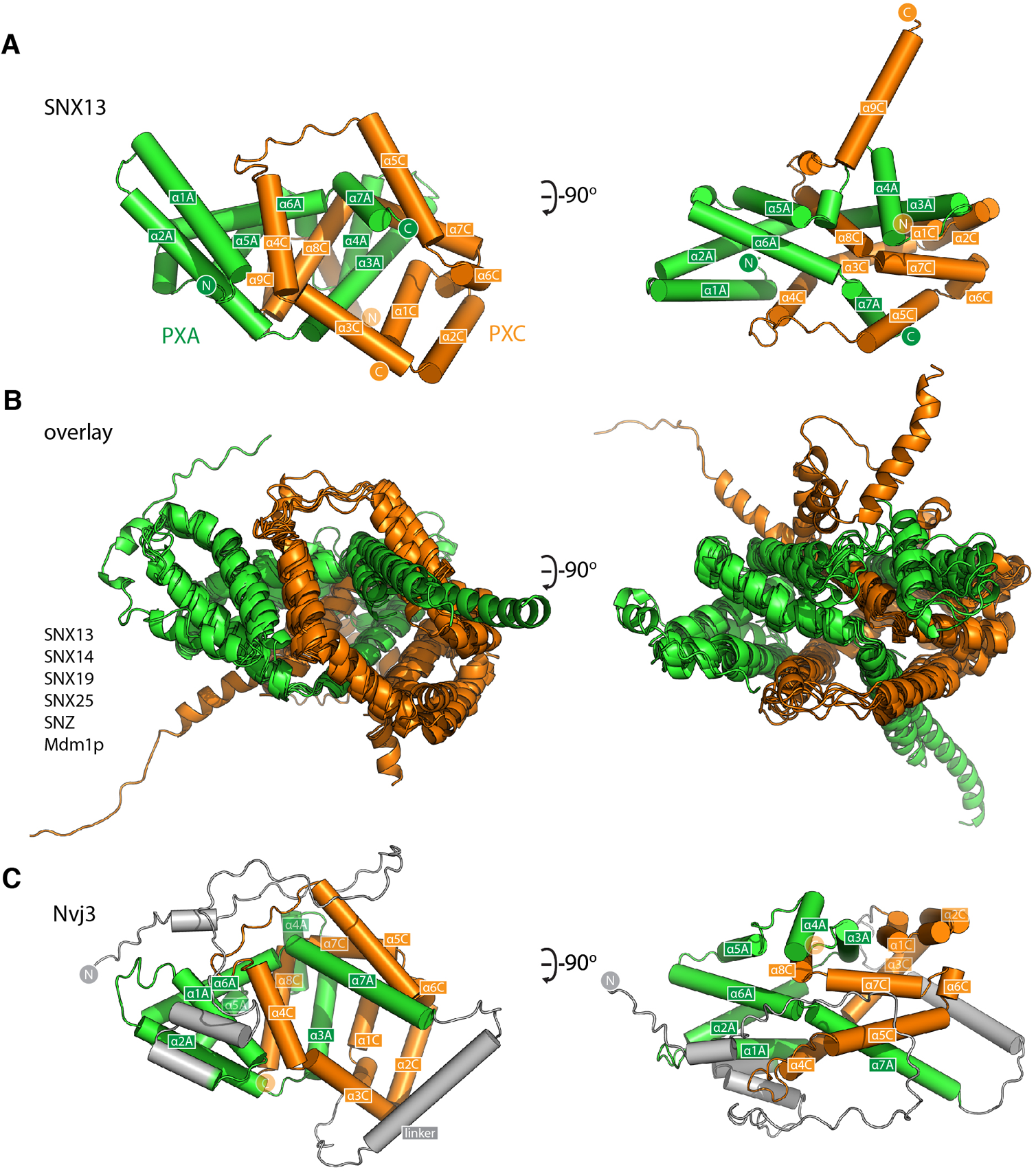
The PXA and PXC domains combine to form an intertwined α-helical structure. (**A**) Structure of the PXA and PXC domains of human SNX13 predicted by AlphaFold2 in green and orange respectively, shown with cylinders for α-helices. The two domains are predicted to be tightly interwoven. (**B**) An overlay of the core PXA-PXC domains of human, yeast and fly SNX-RGS proteins shows that all predicted structures have the same topology. (**C**) Predicted structure of *S. cerevisiae* Nvj3 with the regions expected to be similar to PXA and PXC domains coloured green and orange and the linker shown in grey.

A visual inspection of the core PXA-PXC domains reveals the presence of a large channel that runs through much of the structure with an exposed surface composed of side chains of both domains (**Fig. 4**; **Fig. S5**; **Movie S1**). The channels are extremely hydrophobic, as seen from mapping surface hydrophobicity (**Fig. 4A**; **Fig. S5**) or from inspecting the details of the side chains lining the channel of human SNX13 as an example (**Fig. 4B**; **Fig. S5**). The entrance to the cavities is highly conserved indicating it is likely to be a critical functional element of the proteins (**Fig. 4A**; **Fig. S5**). The presence of such a hydrophobic channel or pocket is a common feature of the class of proteins known as lipid transfer proteins (LTPs) which can bind, extract and transport hydrophobic lipids and fatty acids from various compartments for non-vesicular trafficking ^31–34^ (**Fig. S5**). Given this structural similarity, coupled with the known localisation of the SNX-RGS proteins at ER membrane contact sites and their effects on lipid homeostasis, we propose that this channel in the SNX-RGS proteins is very likely to be a lipid binding pocket. The sizes of the cavities found in the various PXA-PXC proteins are relatively large, ranging from ~2000-6000 Å^3^ compared with typical sized lipid binding pockets in other LTPs of ~400-2000 Å^3^ (**Fig. S5**; **Table S3**). These cavities are larger than those found in cholesteryl ester transfer protein (CETP), which can bind up to two phospholipids and two neutral lipids such as cholesteryl esters or triglycerides at once (**Fig. S5**)^35^. Manual placement of a phosphatidylethanolamine lipid (volume ~1200 Å^3^ for palmitoyl-oleoyl-phosphatidylethanolamine^36^) in the channel of SNX13 shows that the dimensions of this hydrophobic pocket in the SNX-RGS proteins is well suited for binding lipid acyl chains (**Fig. 4C**; **Movie S1**).

**Figure 4.**
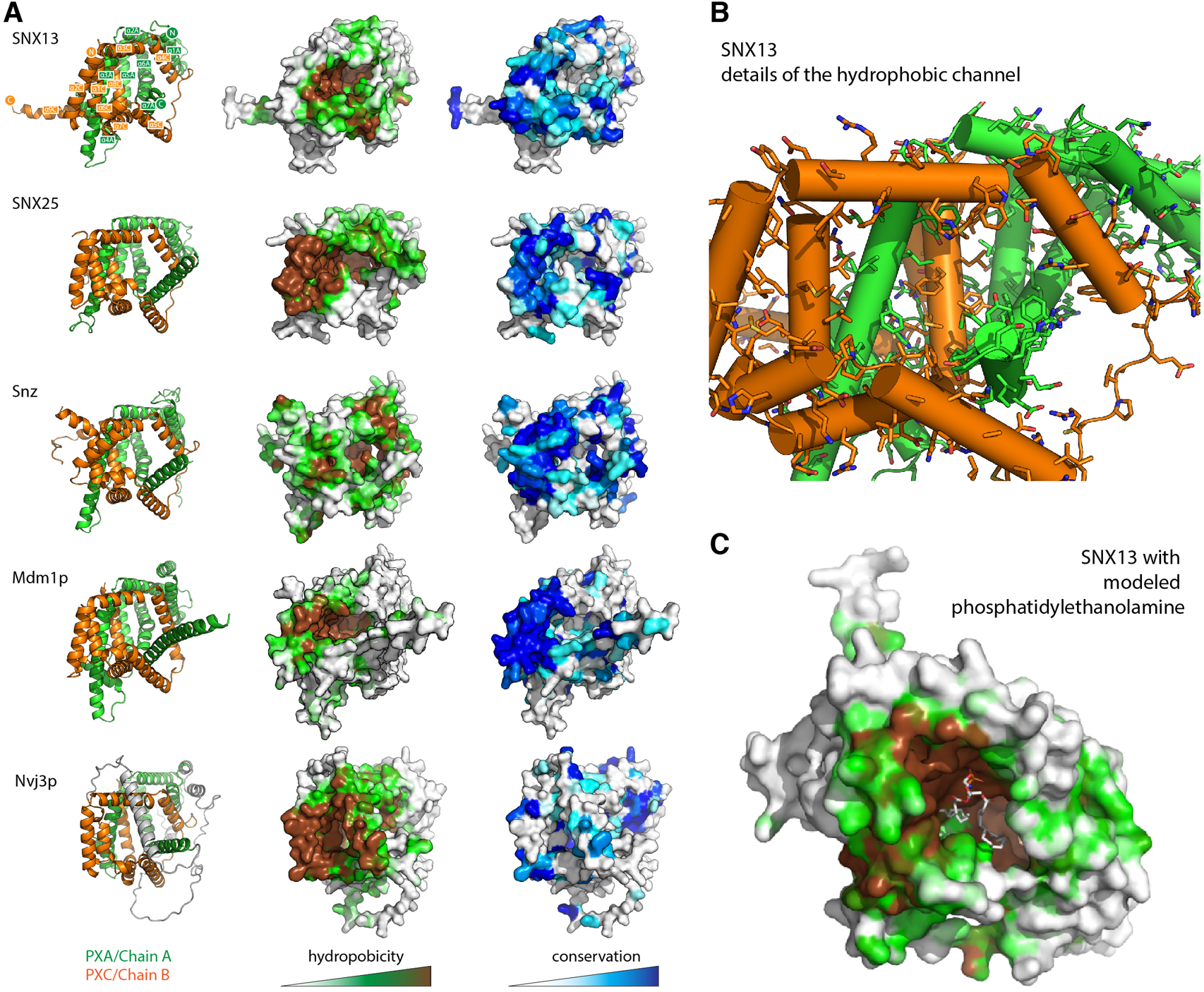
The PXA-PXC structure forms a conserved hydrophobic cavity with potential for lipid binding. (**A**) The predicted structures of PXA-PXC domains from human SNX13 and SNX25, fly Snz, and yeast Mdm1 and Nvj3 are shown in ribbon representation (left), surface coloured by hydrophobicity (middle), and sequence conservation (right). All PXA-PXC structures have a highly conserved hydrophobic tunnel. (**B**) Structure of PXA and PXC domains of SNX13 with sidechains of the putative lipid binding pocket shown. (**C**) The PXA and PXC domains of SNX13 is shown in close-up with its surface coloured for hydrophobicity as in (**A**). A phosphatidylethanolamine lipid has been docked manually in the putative lipid binding pocket to give perspective on the dimensions of the channel.

Several previous studies have shown that the SNX-RGS proteins are rapidly recruited to the surface of newly formed lipid droplets. This has been observed for Mdm1 in yeast ^11^, Snz in flies ^10^, and SNX13, SNX14, and SNX19 in mammalian cells ^6, 7, 9, 13^. This provides strong support for a role in direct transfer of neutral lipids, phospholipids, or lipid metabolites at ER-LD contact sites. We confirmed the ability to localise to the LD surface is conserved in all human family members by analysing the localisation of SNX25 (**Fig. 5**). In A431 cells in the absence of stimulation SNX25 shows a relatively diffuse localisation, however on addition of oleic acid SNX25-FLAG is rapidly redistributed and shows a strong degree of localisation to the surface of newly formed LDs. Further work will be needed to examine if these ER-associated LDs are also in contact with other membrane compartments, but this data confirms that all SNX-RGS family members from yeast to humans have a strong affinity for the LD monolayer.

**Figure 5.**
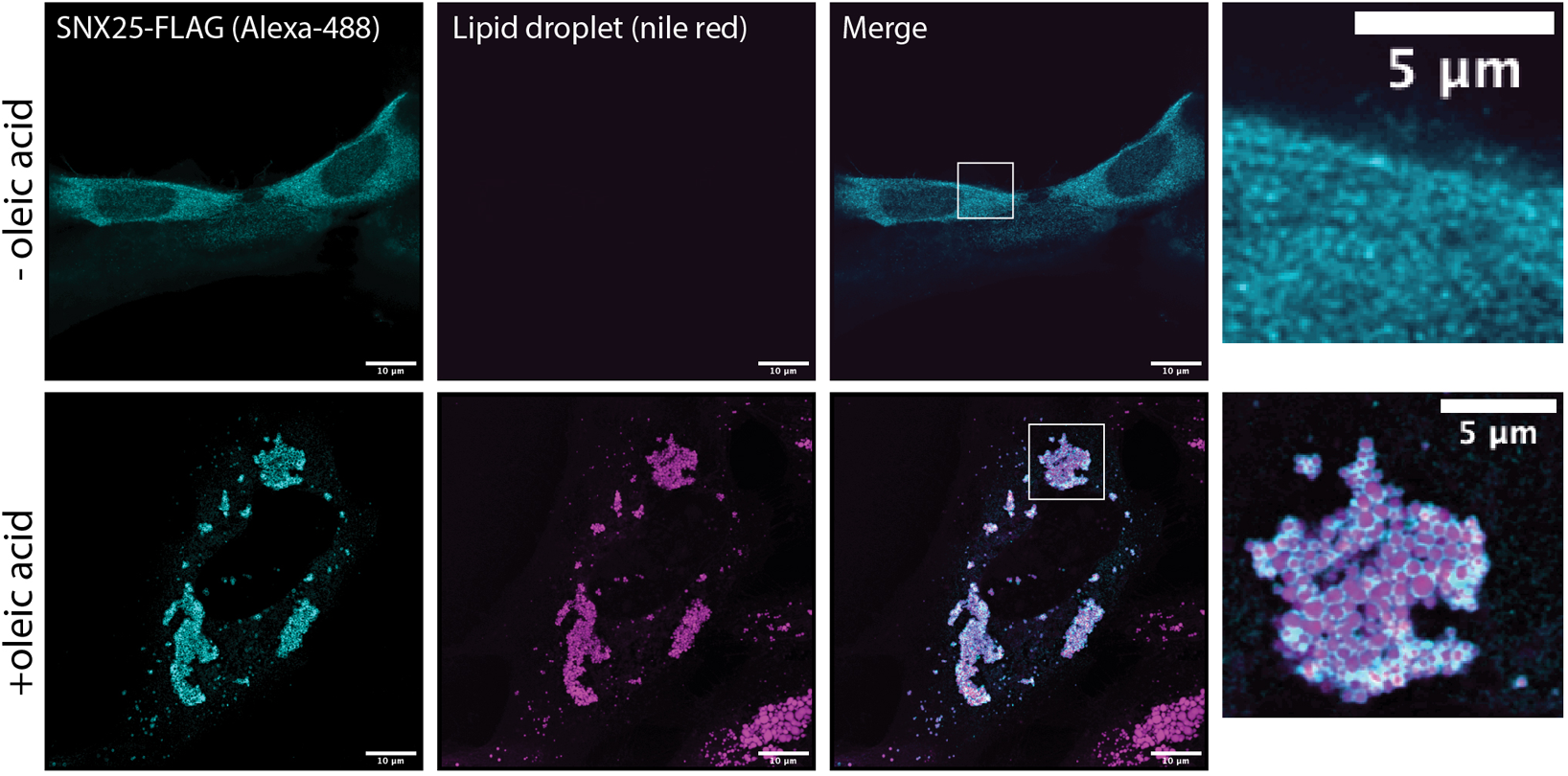
Human SNX25 is localises to newly synthesised LDs following oleic acid addition. A431 cells were transfected with SNX25-FLAG and either left untreated or treated with 50 μg/ml oleic acid to stimulate LD formation. After oleic acid addition SNX25-FLAG undergoes rapid redistribution to the periphery of newly generated LDs LD periphery. Scale bars represent 10 μm.

### Structural bioinformatics of the SNX-RGS proteins

The DALI webserver ^37^ is now capable of searching the entire database of AlphaFold2 predictions ^26^ for structurally similar proteins. We performed a search using the core PXA-PXC structure of SNX25 to determine of any other proteins share similar predicted folds (**Table S2**). This revealed several interesting observations. Firstly, while the PXA-PXC structure is found in proteins across metazoans, plants and yeast we did not find any similar structures in either *D. dictyostelium* or *P. falciparum* and they therefore do not seem to be present in all eukaryotes. Secondly, we identified three proteins in the fission yeast *S. pombe* with this fold. Meiotically up-regulated gene 122 protein (Mug122) (O74444) lacks the RGS domain but has both transmembrane and central PX domains, while snx12 (Q9USN1) has each of the domains common to the SNX-RGS family. The protein annotated as Pxa1 (O14200) appears to be most similar to Nvj3, as it lacks transmembrane, PX and RGS domains. We also performed a BLAST sequence search of the *Chaetomium thermophilum* genome, a thermophilic filamentous yeast popular with structural biologists because of the generally higher stability of the encoded proteins. The *C. thermophilum* genome encodes three proteins with putative PXA-PXC structures (**Table S2**), and these were confirmed by performing ColabFold predictions of their structures (not shown). One of these has all domains common to the SNX-RGS family, a second lacks the RGS domain like human SNX19, and the third has only the PXA-PXC domains like *S. cerevisiae* Nvj3.

### Lec1/Ypr097w is another putative LTP found in yeast and ameoba

While examining the AlphaFold2 predicted structures of PX domain containing proteins in various species, primed by the discovery of the PXA-PXC lipid-binding fold, we noticed that another yeast protein called Lec1/Ypr097w (Q06839) also possessed a superficially similar architecture. Lec1/Ypr097w is 1073 residues in length and is annotated to have a central PX domain between an N-terminal PXB domain and C-terminal domain of unknown function (DUF3818) (**Fig. 6A**). Analogously to the SNX-RGS proteins, the N- and C-terminal domains of Lec1/Ypr097w form α-helical structures with previously unknown folds that come together to form an intimately interwoven structure (**Fig. 6B**; **Fig. S6A and S6B**). Despite this superficial similarity however, neither domain is related in sequence or structure to the PXA or PXC domains and the way they interact with each other is also entirely different. Note that the term ‘PXB domain’ is also used for a C-terminal domain found in human SNX20 and SNX21 which is also completely unrelated in sequence or structure ^38^. For clarity herein we will refer to the N-terminal Lec1/Ypr097w PXB domain as ‘PXYn’ and the C-terminal DUF3818 domain as ‘PXYc’, where PXY denotes ‘PX-associated domains from yeast’.

**Figure 6.**
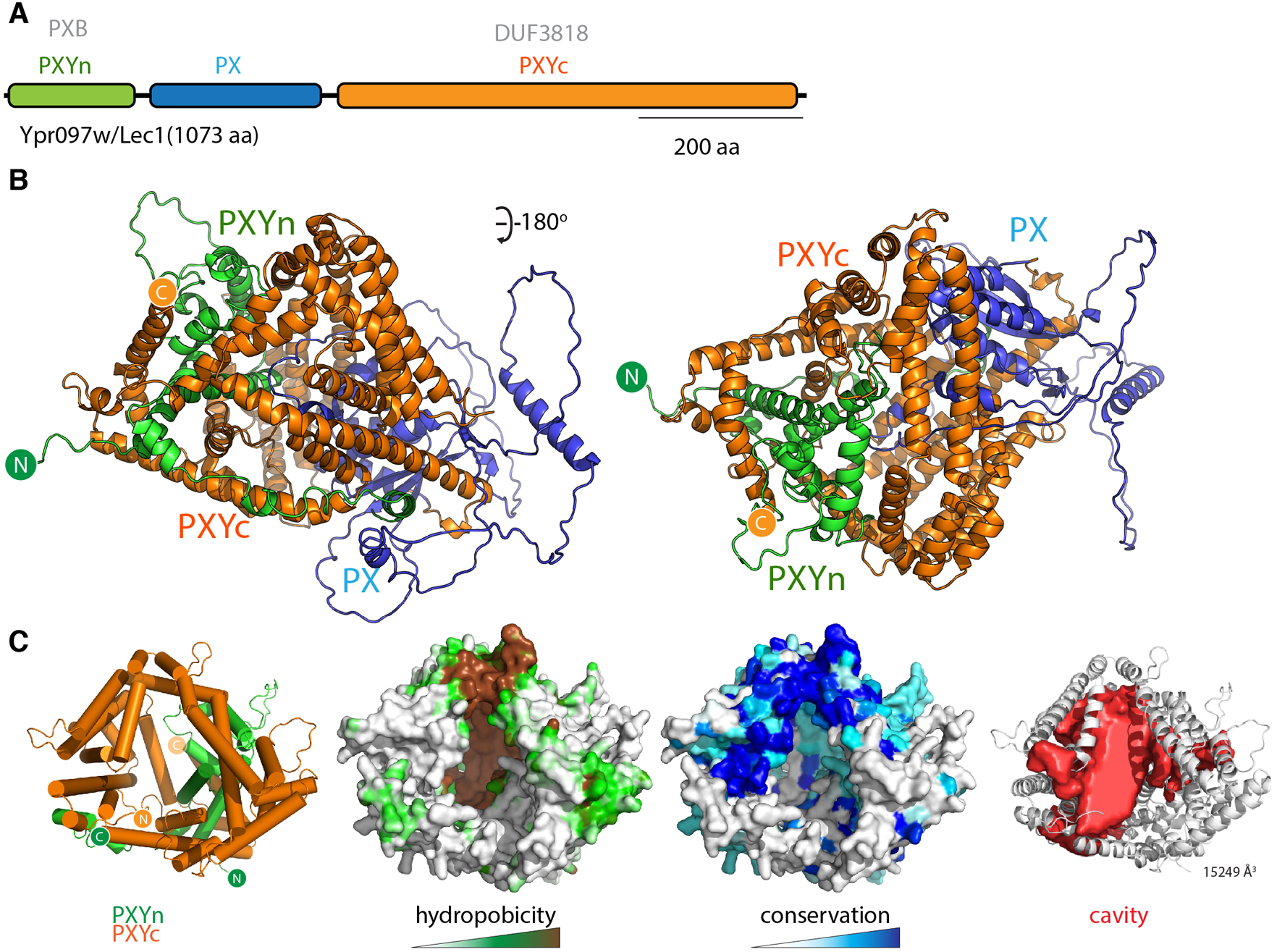
The yeast Lec1/Ypr097w protein forms a unique structure with a hydrophobic cavity. (**A**) Domain structure of the *S. cerevisiae* Lec1/Ypr097w protein. (**B**) AlphaFold2 structural prediction of the Lec1/Ypr097w protein with the PXYn domain in green, PX domain in blue and PXYc domain in orange. (**C**) The PXYn and PXYc domains are shown with α-helices in cylinder representation and the PX domain removed for clarity. The domains encompass a large conserved hydrophobic cavity and like the SNX-RGS proteins may be a potential lipid binding protein. The far right panel shows the solvent accessible cavity (red surface representation) identified with POCASA^56^ and the volume (Å^3^) of the largest identified cavity is indicated.

Most interestingly, the PXYn and PXYc domains of Lec1/Ypr097w also form a barrel-like structure that has a very large cavity surrounded by hydrophobic residues (**Fig. 6C**; **Table S3**). The structure is conserved in homologues from *Schizosaccharomyces pombe* (Q9Y7N9), and *Candida albicans* (A0A1D8PQ52) (**Fig. S6C**). Although structurally unrelated to the SNX-RGS proteins, the general similarities to Lec1/Ypr097w and its homologues are striking, with a central phosphoinositide binding PX domain flanked by N-terminal and C-terminal α-helical domains that come together to form a co-folded intramolecular structure. Like the SNX-RGS proteins based on its structural properties it is very likely that the hydrophobic cavity formed by the PXYn-PXYc domains is also involved in binding to lipids or fatty acids. The volume of the cavity in Lec1/Ypr097w is very large (>15000 Å^3^) and could potentially accommodate multiple phospholipid or neutral lipid molecules. In each of the predicted PXY protein structures from different species the PX domain forms a loose plug over the large hydrophobic cavity suggesting a potential regulatory role for this domain in lipid binding (**Fig. S6C**). This may be relatively dynamic as several independent predictions of Lec1/Ypr097w using ColabFold suggest the PX domain can adopt alternative orientations that either block the pocket or leave the cavity exposed (**Movie S2**).

The PXYn-PXYc structure appears to be relatively restricted throughout evolution, as we only found examples in yeast, including two proteins in *S. pombe*, as well as a protein in *D. dictyostelium. S. pombe* PX domain-containing protein C1450.12 (Q9Y7N9) is highly akin to Lec1/Ypr097w from *S. cerevisiae,* while uncharacterized protein C663.15c (O74521) appears to lack a central PX domain between the two halves of the PXYn-PXYc structure (**Fig. S6**; **Fig. S7**). The *D. discoideum* DUF3818 domain-containing protein (Q54JB8) also lacks a PX domain, and while its C-terminal PXYc domain is like Lec1/Ypr197w, its N-terminal region is structurally divergent from the PXYn domain (**Fig. S7**). Despite this it also possesses a predicted large hydrophobic cavity and therefore may share a functional relationship. Again, we performed a BLAST search of the *C. thermophilum* protein sequences using both *S. pombe* sequences as inputs and found one protein homologous to the Lec1/Ypr097w-like Q9Y7N9 (G0SCP5) and a protein lacking the PX domain like O74521 (G0S997). These discoveries further demonstrate the utility of AlphaFold2 for predicting novel structures, and its power in aiding evolutionary analyses of protein homology.

## Discussion

With the recent advances made by the AlphaFold2 machine learning algorithm and public release of the AlphaFold2 database ^25, 26^, it is now possible to perform detailed structural analyses and evolutionary comparisons of protein families across diverse species. In this work we have solved the crystal structure of the human SNX25 RGS domain and gone on to examine the previously unexplored structures of the full-length SNX-RGS proteins. Although the overall structures of these proteins display conformational flexibility with respect to their N-terminal transmembrane ER tethers, in contrast to previous pictures based on primary and secondary structure predictions the disparate N- and C-terminal PXA and PXC domains form a highly entwined intramolecular interaction generating a compact cylindrical α-helical structure. The most notable feature of a central hydrophobic cavity with a highly conserved entrance and surface lining. This leads us to conclude that the likely function of the PXA-PXC structure will be to bind and transport lipids or fatty acids, and that the proteins represent a distinct class of ER membrane-anchored LTPs.

Broader analysis of other PX domain-containing protein family members indicate that the yeast protein Lec1/Ypr097w and related molecules share a superficially similar LTP structure and function. Here the central PX domain is found between N-terminal PXYn and C-terminal PXYc domains that also become conjoined to form an all α-helical barrel-like structure with a large hydrophobic cavity, although their sequences are not related and their structural details are entirely different. As we were performing these studies, a preprint was published by the Schuldiner lab that identified novel proteins that reside at or regulate formation of membrane contact sites in yeast ^24^. In this work Ypr097w was identified to localise to contacts between lipid droplets and the cell surface and renamed Lec1 (Lipid Droplet Ergosterol Cortex 1) because of a potential role in ergosterol transport. This team also analysed the AlphaFold2 predicted structure of Lec1/Ypr097w and came to similar conclusions about its potential role as a novel lipid transfer protein (LTP), as well as identifying the superficial structural similarities to the SNX-RGS proteins.

What do these new structural insights suggest about the function(s) of the SNX-RGS proteins? The hydrophobic nature and conserved surface of the large interior cavity found in the PXA-PXC domain structure indicates they are very likely to mediate the binding of lipids, fatty acids or other lipid-derived metabolites. A general finding is that the SNX-RGS proteins reside in the ER at steady state via their N-terminal membrane anchor, and subsequently colocalise with LDs particularly under conditions of new LD synthesis. They are typically found to be enriched in regions of the ER and ER-associated LDs that are in contact with other membranes where they can play a tethering role via their phosphoinositide-binding PX domain. These membrane contacts include the cell surface in the fat body cells of fruit flies ^10^, the nuclear-vacuolar junction in budding yeast ^8, 11, 12^, and with endolysosomal compartments in mammalian cells ^6, 7, 9, 13^. This localisation to ER-LD-membrane contact sites is dramatically altered under conditions of lipid flux (e.g. addition of excess free fatty acids to cells ^7, 9^), and the SNX-RGS proteins play important roles in lipid metabolism and homeostasis.

The effects of mutations, knockouts or overexpression of SNX-RGS proteins are complex, but at the cellular level their depletion commonly leads to higher cellular levels of neutral triacylglycerol (TAG) and accumulation of cholesterol in late endosomal compartments ^6, 11, 13, 16^. Yeast Mdm1 and human SNX14 have been associated with Faa1 and ACSL3 respectively, long-chain-fatty-acid--CoA ligases that activate fatty acids for synthesis of cellular lipids and degradation via β-oxidation ^7, 11^. Drosophila Snz and human SNX14 were also found to associate with stearoyl-CoA desaturases Desat1 and SCD1 respectively, which catalyze the insertion of a cis double bond at the D-9 position of fatty acyl-CoA substrates ^10, 23^. Knockout of SNX14 causes cells to be sensitive to saturated fatty acid (SFA)-induced toxicity and increased levels of SFAs are incorporated into phospholipids, which can be rescued by SCD1 overexpression. A recent CRISPR screen for genes that influence cholesterol homeostasis found SNX13 (and SNX14) as a prominent hit ^13^. Niemann Pick type C (NPC) disease is caused by genetic defects in the lysosomal cholesterol transport system, Niemann Pick C1 and C2 proteins (NPC1 and NPC2) leading to endolysosomal accumulation of cholesterol and Bis(monoacylglycero)phosphate (BMP, also known as LBPA) ^39, 40^, which is essential for normal cholesterol transport out of the endolysosomal compartment ^41–45^. In screens using the NPC1 inhibitor U18666A cholesterol accumulation could be reversed by depletion of SNX13, and cholesterol was exported in an NPC1-independent manner to other compartments including the plasma membrane. Surprisingly SNX13 depletion led to a significant increase in total lysosomal BMP, which may be important for the observed NPC1-independent cholesterol transport ^46, 47^. Although highly speculative one possibility is that the PXA-PXC domains are important for transport and regulation of lipids such as BMP at contacts between the ER and endolysosomal compartments. Identifying specific ligand(s) and whether the different family members have alternative binding activities will require further experimentation.

The PXA-PXC domains were previously considered as conserved but essentially independent structures in the SNX-RGS family of proteins. From the analyses presented here it is clear that these two domains are intimately associated in a single structural unit and that neither the PXA or PXC sequences are likely to fold correctly in the absence of the other. Previously, several studies have examined the impact of deleting these regions on SNX-RGS protein function. While deletion of either domain generally renders the proteins non-functional as expected, there is evidence that the two sequences contain regions that are important for LD targeting and potentially protein interactions. In yeast, it was found that the PXA domain of Mdm1 could define sites of LD formation, and could recruit the Faa1 fatty-acid—CoA ligase ^12^. In contrast, several reports have shown that the PXC domain by itself is required and sufficient for targeting to the surface of newly formed LDs in both flies and mammals ^7, 10, 13^. Deleting an amphipathic helical region in SNX14 PXC (residues 801-819), which is predicted to sit at the entrance of the hydrophobic cavity of the assembled PXA-PXC structure (**Fig. S5**), prevents LD attachment. These studies suggest that even in the absence of the PXA domain, and thus proper folding of the PXA-PXC structure, there are sequences in the PXC domain capable of direct association with the LD monolayer.

In conclusion, although further experiments are required to experimentally validate these models and define the functional lipid interactions of the SNX-RGS proteins and their distantly related Lec1/Ypr097w cousins, these structural studies provide strong evidence for a novel lipid transport activity mediated by these conserved molecules. As a final general observation, this work further confirms the ability of AlphaFold2 to predict not only already known folds, but to define completely new structures without previously recognised topologies and thus provide new insights into biological function.

## Materials and methods

### Cloning, expression purification of the SNX25 RGS domain

The genes encoding the human SNX25 RGS domain (residues 446-569; A0A494C0S0) and the SNX25 RGS(C526A) mutant were synthesised by Gene Universal (USA) and cloned into the expression plasmid of pGEX-4T-2 for bacterial expression with an N-terminal GST tag and thrombin cleavage site. was construct was transformed into *Escherichia coli* BL21-CodonPlus (DE3)-RIL cells and plated on lysogeny-broth (LB) agar plates supplemented with ampicillin (0.1 mg/mL). Single colonies were then used to inoculate 50 mL of LB medium containing ampicillin and the culture was grown overnight at 37°C with shaking at 180 rpm. The following day 1 L of LB medium containing ampicillin (0.1 mg/mL) was inoculated using 10 ml of the overnight culture. Cells were then grown at 37°C with shaking at 200 rpm to an optical density of 0.8-0.9 at 600 nm and the protein expression was induced by adding 0.5 mM IPTG (isopropyl-β-D-thiogalactopyranoside). Expression cultures were incubated at 20°C overnight with shaking and the cells were harvested the next day by centrifugation at 4000 rpm for 15 min using a Beckman rotor JLA 8.100. Cell pellets were resuspended in 20 mL (for cell pellet from 1 L culture) of lysis buffer (50 mM HEPES, (pH 7.5), 500 mM NaCl, 5% glycerol, Benzamidine (0.1 mg/mL), and DNAse (0.1 mg/mL)). Resuspended cells were lysed by using the cell disrupter (Constant systems, LTD, UK, TS-Series) and the soluble fraction containing the protein was separated from cell debris by centrifugation at 18,000 rpm for 30 min at 4°C. The soluble fraction was first purified by affinity chromatography using Glutathione Sepharose 4B resin (GE Healthcare) and the GST tag was cleaved on column by incubating the protein with Thrombin (Sigma Aldrich) overnight at 4°C. The next day the protein was eluted using 50 mM HEPES (pH 7.5), 200 mM NaCl. The eluted protein was then concentrated and further purified by gel filtration chromatography (Superdex 75 (16/600), GE Healthcare) using 50 mM HEPES, pH 7.5, 200 mM NaCl, 0.5 mM TCEP (tri(2-carboxyethyl)phosphine) and the fractions corresponding to SNX25 RGS domains were analysed by SDS PAGE.

### Crystallisation and structure determination of the SNX25 RGS domain

Purified proteins were concentrated to 12 mg/mL for crystallisation. Initially 96 well crystallisation screens were set up using the hanging drop method with a Mosquito Liquid Handling robot (TTP LabTech). The optimised diffraction-quality crystals were obtained in 24-well plates using hanging drop method. The SNX25 RGS domain was crystallised in 30% PEG 4000, 0.2 M Sodium acetate, 0.1 M Tris pH 8.5, while the SNX25 RGS(C526A) domain was crystallised in 18% PEG 8000, 0.1 M calcium acetate, 0.1 M sodium cacodylate, pH 6.5. Diffraction data was collected at the Australian Synchrotron MX2 beamline (Clayton, VIC, Australia), integrated with XDS ^48^ and scaled with AIMLESS software ^49^. The SNX25 RGS structure was solved by molecular replacement using the RGS1 structure as input (PDB ID 2BV1; ^28^) in PHASER ^50^ and the resulting model was rebuilt and refined by using COOT ^51^ and PHENIX ^52^ respectively. The SNX25 RGS(C526A) was solved similarly by molecular replacement using the wild-type SNX25 RGS domain as input.

### Protein structural prediction, modelling and visualisation

The structural predictions of the full-length proteins from different species including *S. cerevisiae, D. melanogaster, C. elegans, P. falciparum, D. discoideum, A. thaliana, S. pombe, M. janaschii, D. rerio* and *H. sapiens* analysed in this study were obtained from the AlphaFold2 database (https://alphafold.ebi.ac.uk; ^25, 26^). To generate predicted models of human SNX25 including the previously annotated transmembrane domain ^5, 29^ and the *S. cerevisiae* Lec1/Ypr097w to assess the flexibility of its PX domain, we used the AlphaFold2 neural-network ^25^ implemented within the freely accessible ColabFold pipeline ^30^. For each modelling experiment ColabFold was executed using default settings where multiple sequence alignments were generated with MMseqs2 ^53^ and structural relaxation of final peptide geometry was performed with Amber ^54^ to generate three models per structure. Sequence conservation was mapped onto the modelled SNX27 FERM domain structure with Consurf ^55^. Interior cavities of proteins were mapped and their volumes were calculated using POcket-CAvity Search Application (POCASA)^56^, with default settings of 2 Å for probe radius and 1 Å for the grid size. All structural images were made with Pymol (Schrodinger, USA; https://pymol.org/2/).

### Localisation of SNX25 in A431 cells

The human SNX25 open reading frame (ORF) cloned into the pcDNA3.1+-C DYK was obtained from Genscript (NM_031953). This expresses a C-terminally tagged SNX25 (SNX25-FLAG) that has the same sequence as the AlphaFold2 model shown in **Fig. 2B**. Note however, that it lacks the predicted N-terminal ER anchor found in the SNX25 isoform A0A494C0S0 that is modelled in **Fig. S4**. A431 cells were maintained in DMEM medium (Gibco Life technologies) supplemented with 10% foetal bovine serum (FBS) and Penicillin/Streptomycin. A431 cell line was sourced from ATCC and tested regularly for mycoplasma contamination. Cells (2 × 10^5^) were plated in 6 well culture dishes (Nunc™, Cat. No. 140675, Culture area—9.6 cm^2^) and grown to 50% confluency before transfection of SNX25-FLAG construct using lipofectamine 3000 (Invitrogen) as per manufacturer’s protocol. Cells were shifted to DMEM media containing oleic acid (50 µg/ml) 4 h post transfection and incubated for another 20 h. DMEM with oleic acid was prepared by mixing oleic acid (Sigma Aldrich, Cat. No. O-1008) in DMEM medium containing bovine serum albumin (fatty acid free) to final oleic acid concentration (50 µg/ml) with rigorous shaking at 37°C to avoid precipitation. Cells were fixed 24 h post-transfection with 4% paraformaldehyde in phosphate buffered saline (PBS) at 4°C and subsequently permeabilised with 0.1% Triton X-100 in PBS for 7 min. Cells were probed with FLAG antibody (Sigma Aldrich Cat. No. F7425, 2 µg/ml) and anti-Rabbit secondary antibody Alexa Fluor®488 conjugate (4 µg/ml) to visualise SNX-25 protein. Cells were finally labelled with nile red solution (100 nM) to visualise lipid droplets. Confocal images (1024 × 1024) were acquired on a Zeiss inverted LSM 880 microscope coupled with fast airyscan detector (Carl Zeiss, Inc) equipped with 63X oil immersion objective, NA 1.4.

## Acknowledgements

We thank Prof. Rob Parton for the gift of A431 cells, and antibodies for LD staining. This work was supported by funding from the Australian Research Council (ARC) (DP200102551). BMC is supported by an NHMRC Senior Research Fellowship (APP1136021). WMH is supported by funds from the Welch Foundation (I-1873), the NIH NIGMS (GM119768), NIDDK (DK126887), Ara Parseghian Medical Research Fund, and the UT Southwestern Endowed Scholars Program.

## Author contributions

Conceptualization, WMH, BMC.; Methodology, SW, BP, VT, BMC; Investigation, SW, BP, VT, BMC; Writing – Original Draft, SW, BMC; Writing – Review & Editing, SW, BP, VT, BMC; Funding Acquisition, WMH, BMC; Supervision, WMH, BMC.

## Conflict of interest

Authors declare that they have no conflict of interest.

## Supplementary Information

**Figure S1.**
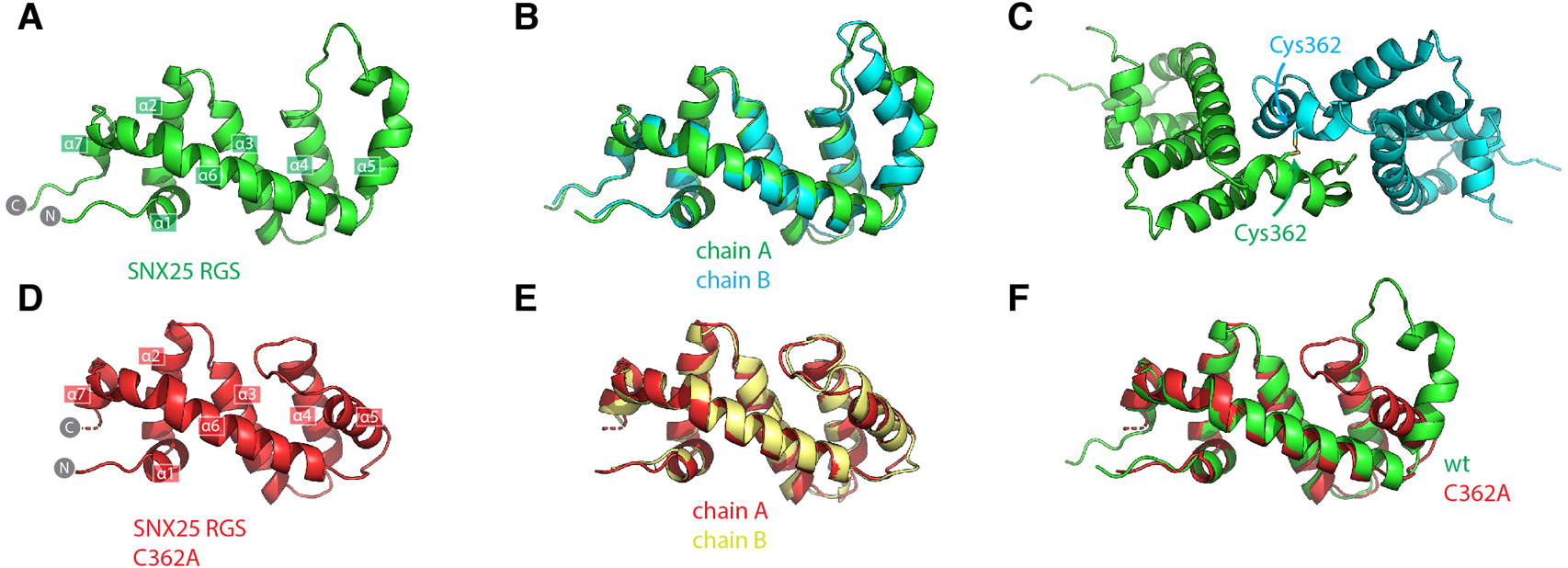
Crystal structure of the SNX25 RGS domain. (**A**) Crystal structure of the wild-type human SNX25 RGS domain. (**B**) Alignment of the two chains in the asymmetric unit of wild-type SNX25 RGS domain. (**C**) The position of the non-native disulfide bond formed between adjacent chains in the crystal lattice. This causes the change in orientation of the α5 helix. (**D**) Crystal structure of the SNX25(C526A) mutant protein. (**E**) Alignment of the two chains in the asymmetric unit of SNX25(C526A) RGS domain. (**F**) Structural alignment of the wild-type and C526A SNX25 RGS domain structures.

**Figure S2.**
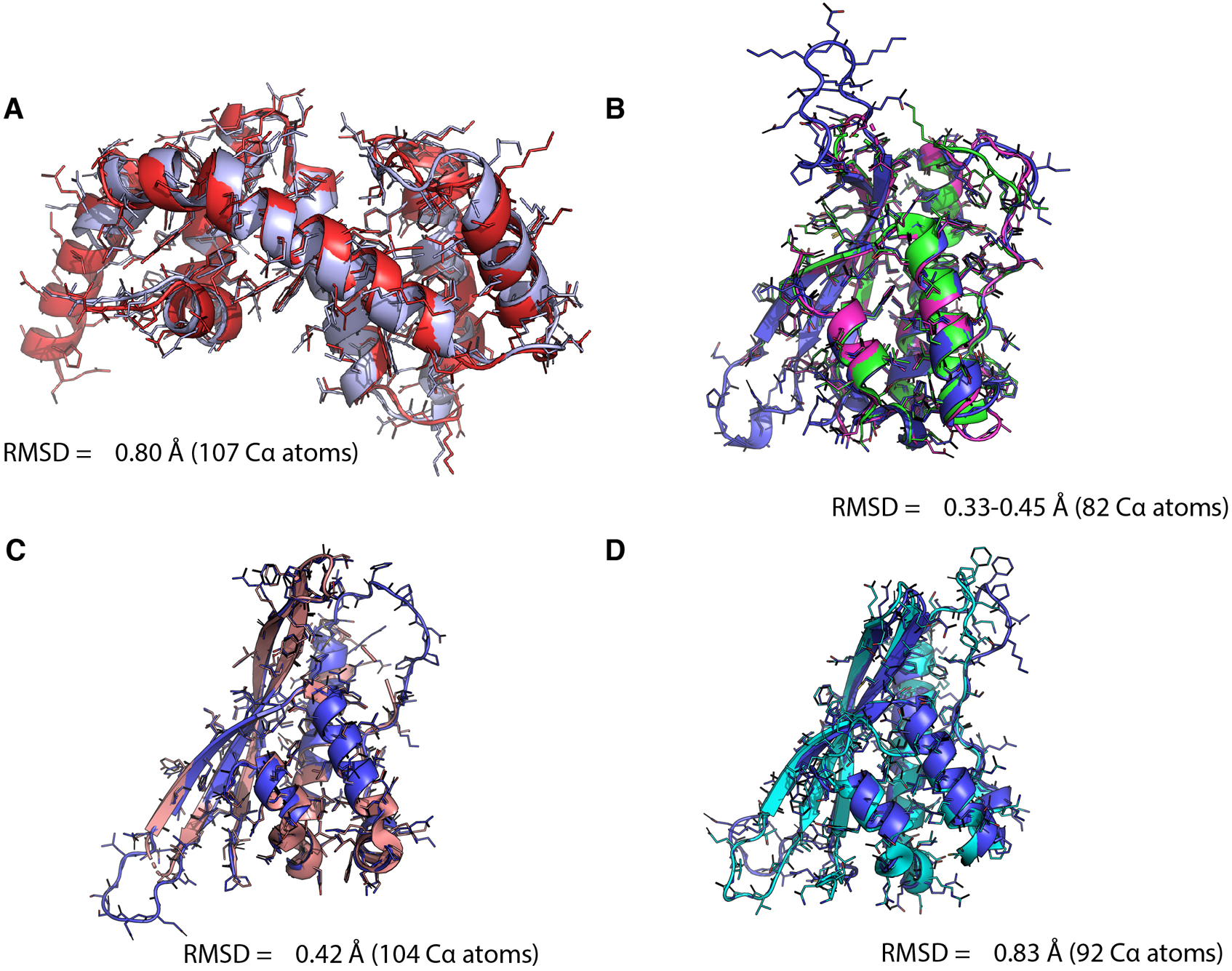
Comparison of experimental SNX-RGS structures with their AlphaFold2 predictions. (**A**) Crystal structure of the human SNX25(C326A) RGS domain (red, this study) aligned with the AlphaFold2 prediction (light blue). (**B**) Crystal structures of human SNX14 PX domain (green, PDB 4PQP; magenta, 4PQO) with the AlphaFold2 prediction (blue). (**C**) Crystal structure of mouse SNX19 PX domain (pink, PDB 4P2I) aligned with the AlphaFold2 prediction (blue). (**D**) NMR structure of human SNX25 PX domain (cyan, PDB 4PQP) aligned with the AlphaFold2 prediction (blue).

**Figure S3.**
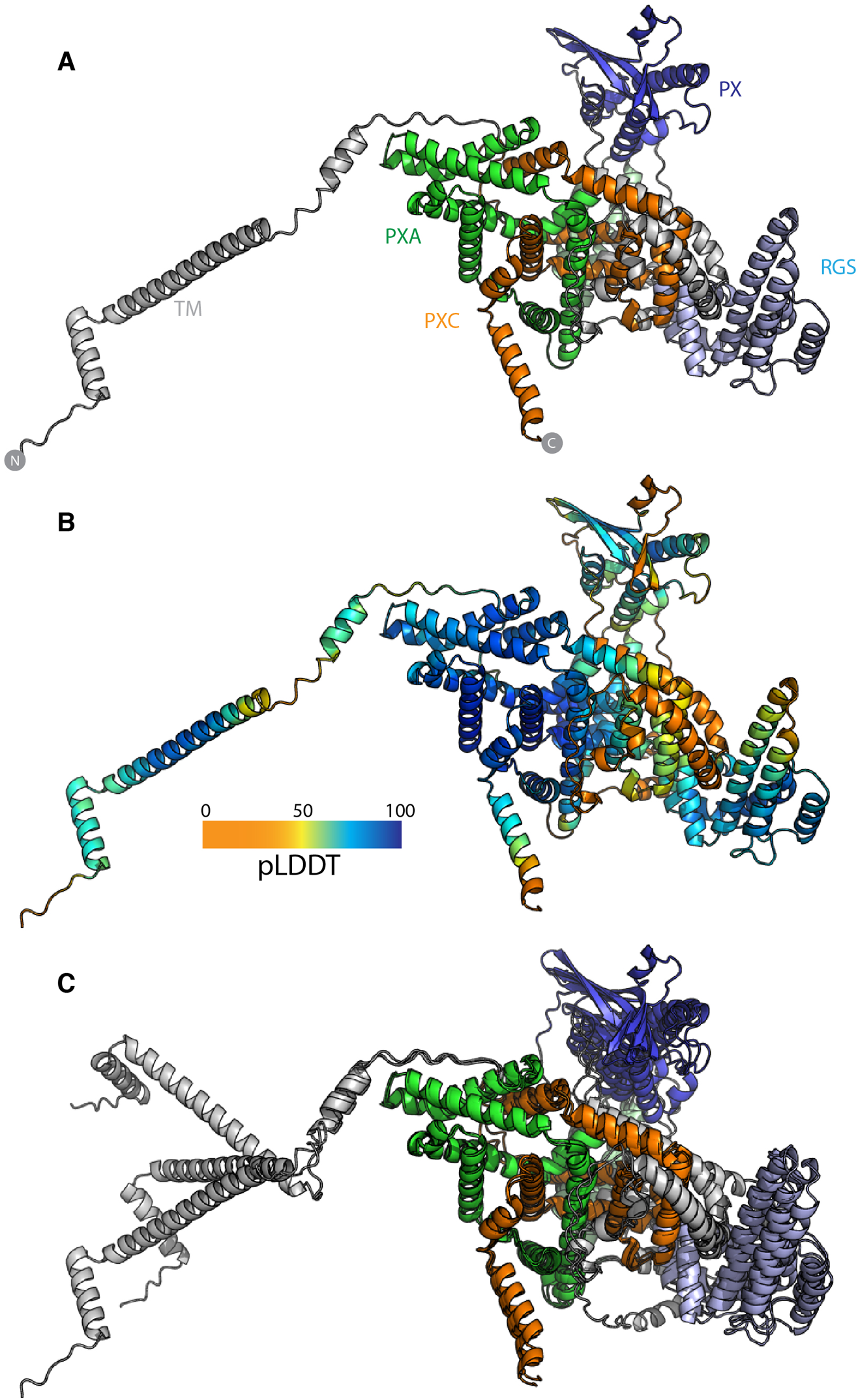
Additional details of the SNX13 predicted full length structure. (**A**) AlphaFold2 prediction of full-length human SNX13 protein with individual domains coloured as in Fig. 2B with the TM domain and linker regions (grey), PXA domain (green), RGS domain (light blue), PX domain (blue), and PXC domain (orange). (**B**) AlphaFold2 prediction of full-length human SNX13 protein coloured according to the pLDDT score. (**C**) The AlphaFold2 predictions of human, mouse and zebrafish SNX13 proteins were aligned based on the core PXA-PXC structure.

**Figure S4.**
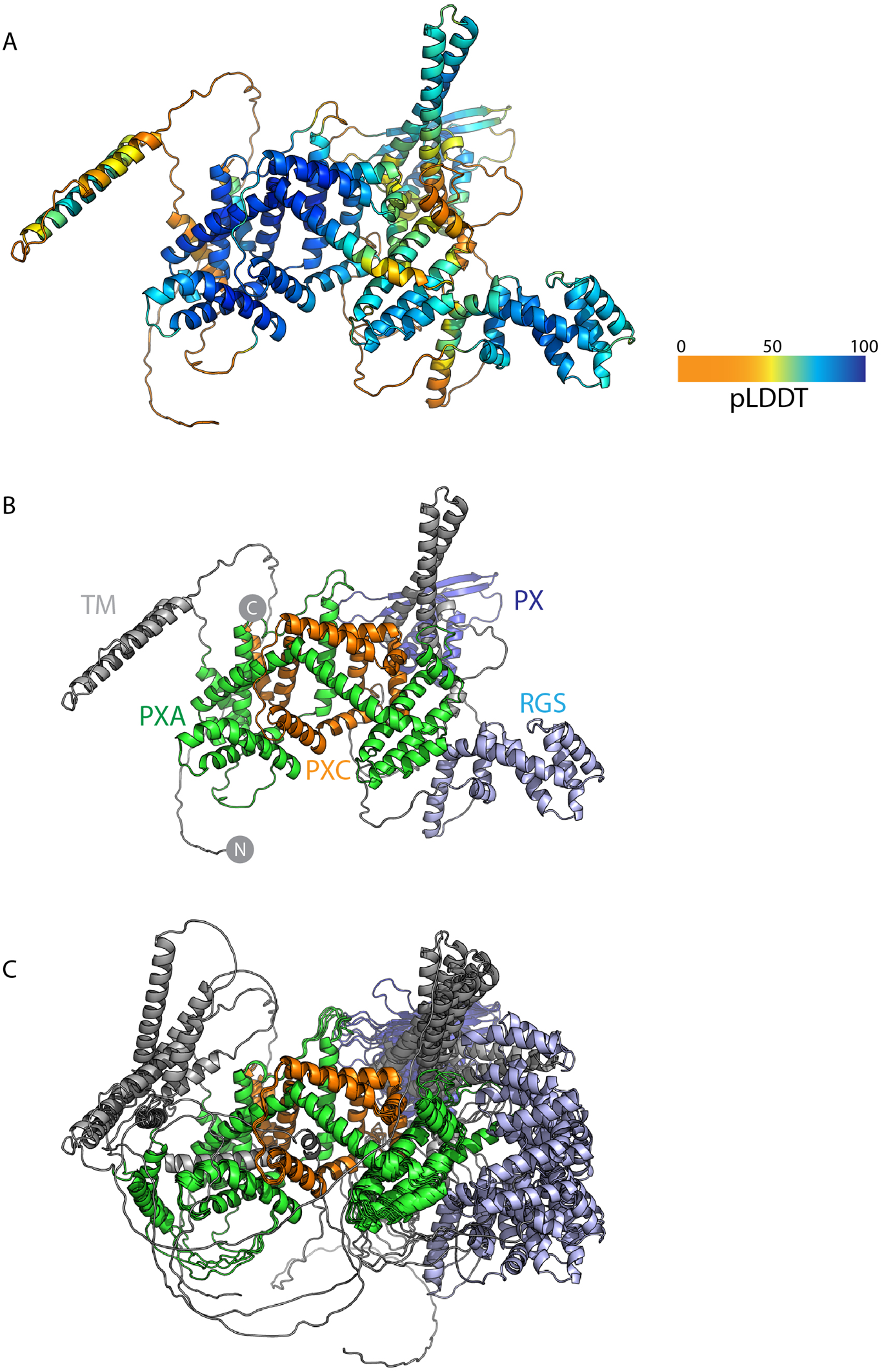
AlphaFold2 prediction of full-length human SNX25 protein. (**A**) Top ranked structure of full length human SNX25 ^5^ (Uniprot ID A0A494C0S0) predicted by ColabFold coloured according to pLDDT score. (**B**) Top ranked structure of human SNX25 predicted by ColabFold coloured according to the indicated domains. (**C**) Alignment based on the core PXA-PXC structure of the five structural predictions of human SNX25 from ColabFold.

**Figure S5.**
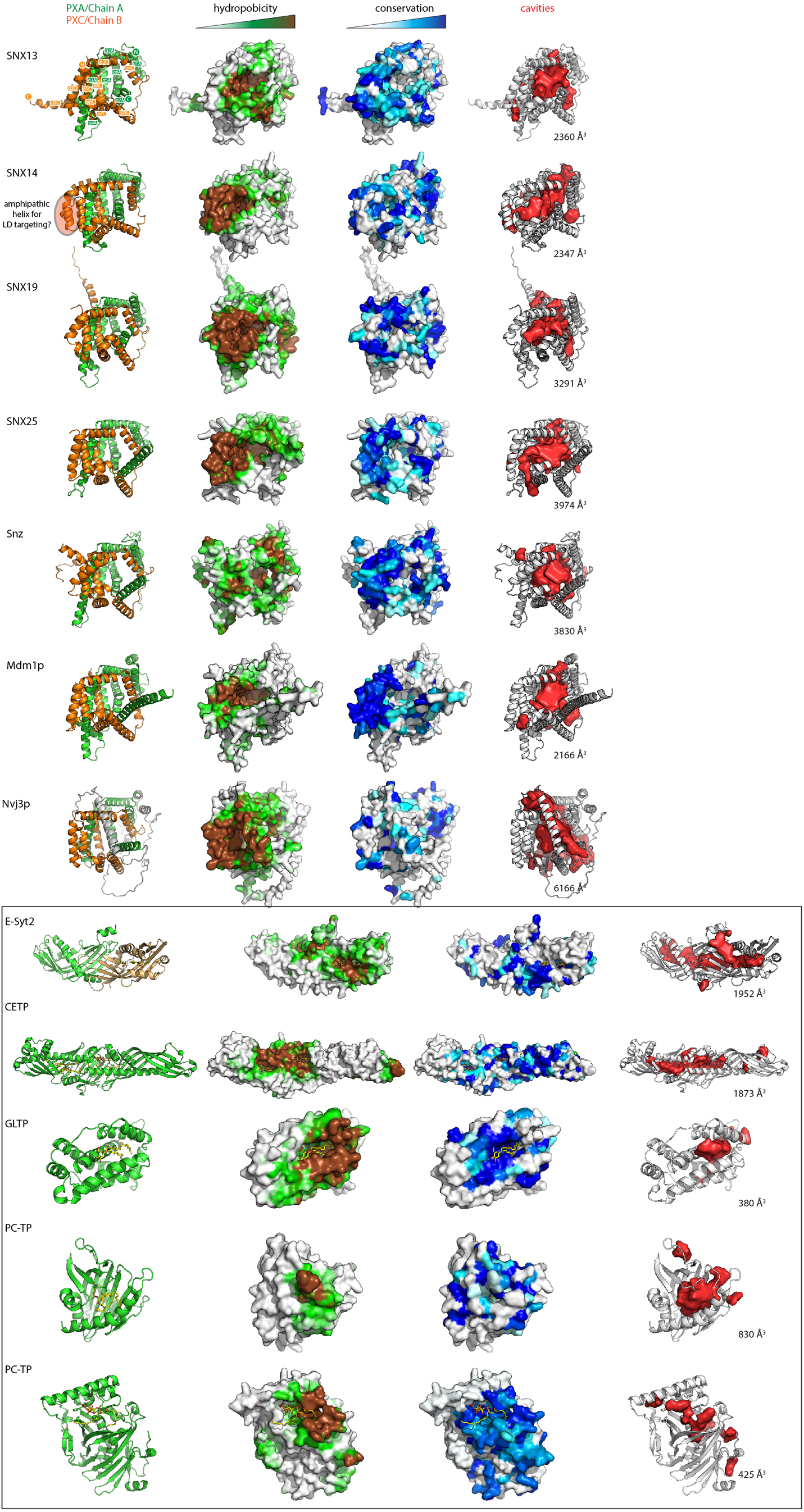
Conserved hydrophobic cavities in all human, yeast and fly PXA-PXC domains. Structures of indicated PXA-PXC domains in ribbon (left) surface coloured by hydrophobicity, and sequence conservation. A region in SNX14 is highlighted that when deleted prevents LD recruitment^7^. Far right panels display accessible cavities (red surface representation) in the proteins identified with POCASA^56^ and the volume (Å^3^) of the largest continuous cavity is indicated. The five bottom structures are examples of other lipid transfer proteins bound to different lipids, demonstrating how similar kinds of conserved hydrophobic pockets can serve as lipid binding cavities and provided for comparison. Extended synaptotagmin 2 (E-Syt2) is in complex with a phosphatidylethanolamine lipid and Triton-X100 detergent molecule (PDB 4P42) ^57^. Cholesteryl ester transfer protein (CETP) is in complex with two cholesteryl esters and two phosphatidylcholine lipids (PDB 2OBP) ^35^. Glycolipid transfer protein (GLTP) is in complex with N-oleoyl-glucosylceramide (PDB 3S0K) ^58^. Phosphatidylcholine transfer protein (PC-TP) is shown in complex with a phosphatidylcholine molecule (PDB 1LN2) ^59^. Human OSBP-related protein 1 (ORP1) is shown bound to phosphatidylinositol(4,5)P_2_ (PDB 5ZM6)^60^

**Figure S6.**
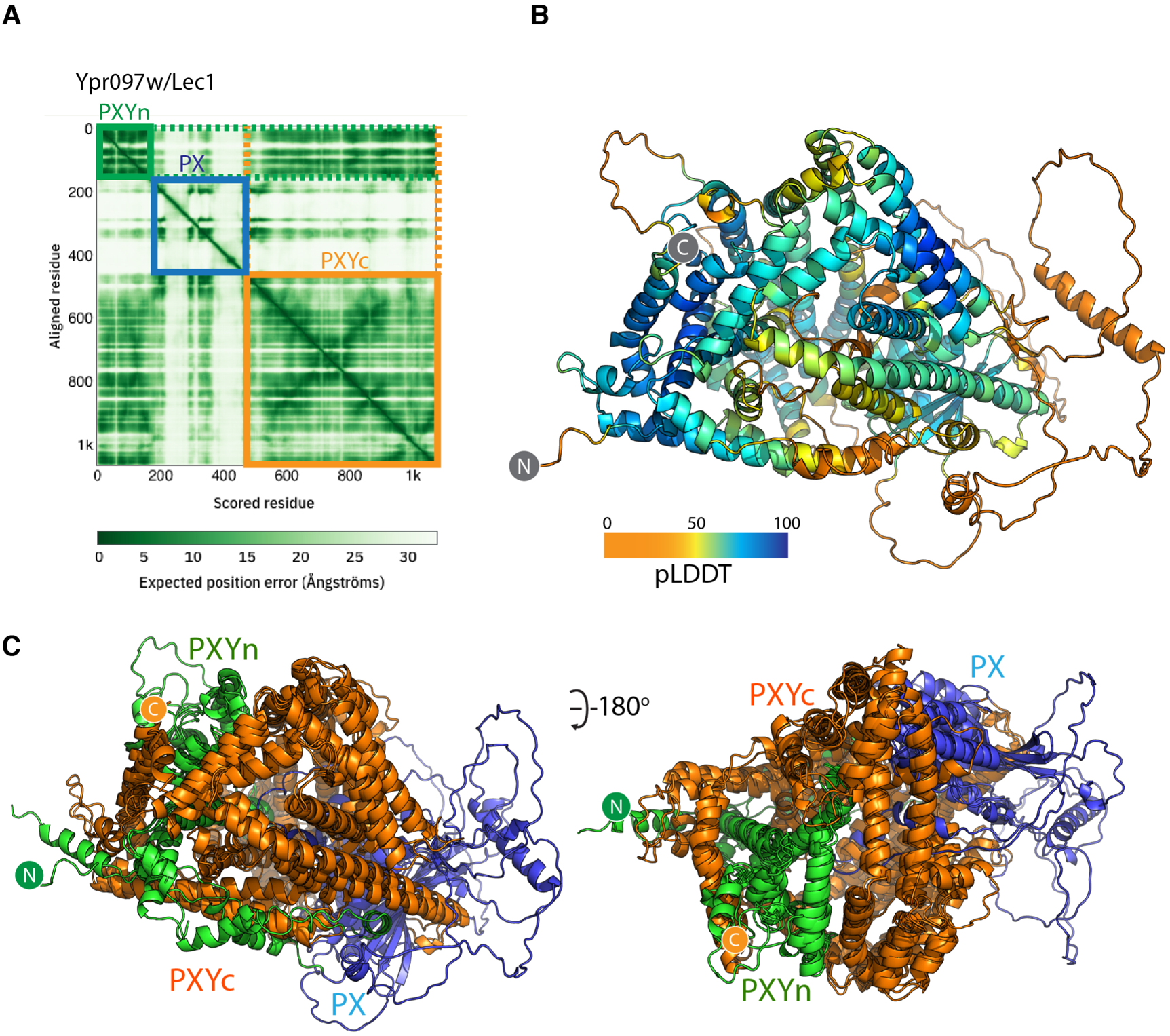
AlphaFold2 predicted structure of Lec1/Ypr097w. (**A**) Plot of the Predicted Alignment Error (PAE) from the AlphaFold2 database. There is a strong degree of correlation between the N-terminal PXYn and C-terminal PXYc domains suggesting these two domains are physically associated. (**B**) The predicted structure of Lec1/Ypr097w from S. cerevisiae coloured according to the pLDDT score. (**C**) Overlay of the PXY proteins from S. cerevisiae, S. pombe and C. albicans.

**Figure S7.**
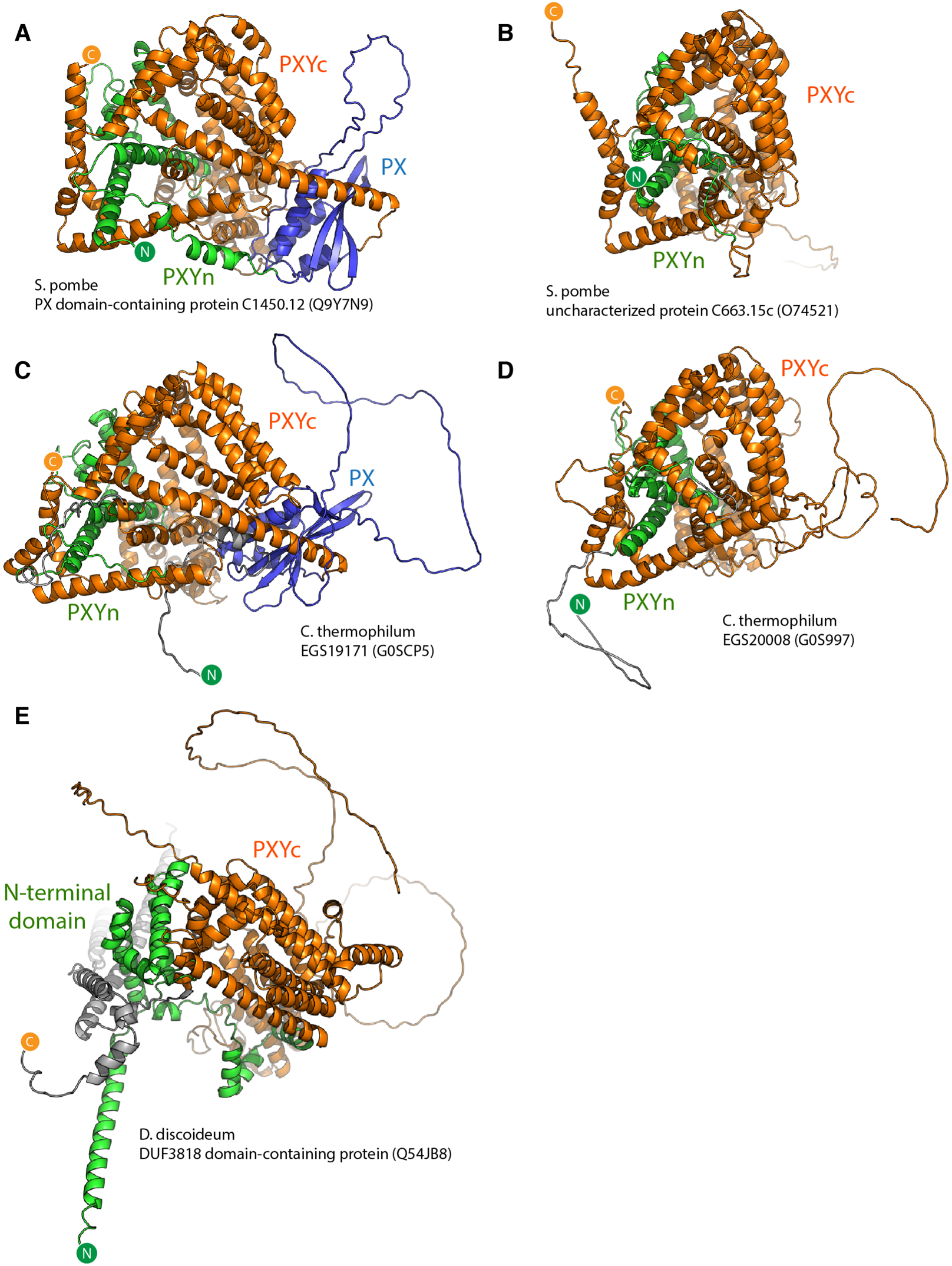
AlphaFold2 predicted structure of PXY domain proteins found in *S. pombe*, *C. thermophilum* and *D. dictyostelium*. (**A**) Structural prediction of *S. pombe* Lec1/Ypr092w orthologue C1450.12. (**B**) Structural prediction of *S. pombe* Lec1/Ypr092w homologue C663.15c. (**C**) Structural prediction of *C. thermophilum* Lec1/Ypr092w orthologue EGS19171. (**D**) Structural prediction of *C. thermophilum* Lec1/Ypr092w homologue EGS20008. (**E**) Structural prediction of *D. discoideum* DUF3818 domain-containing protein.

**Table S1.** Statistics for X-ray crystallographic data collection and structure refinement^a^.

|  | Human SNX25 RGS<br>domain | Human SNX25 RGS<br>domain (C526A) |
| --- | --- | --- |
| **Wavelength (Å)** | 1.006 | 0.95364 |
| **Resolution range (Å)** | 45.84 -2.4 (2.486-2.4) | 48.25-2.42 (2.52-2.42) |
| **Space group** | P 1 21 1 | P 1 21 1 |
| **Unit cell** | 48.6 57.1 67.0 90 109.5<br>90 | 44.0 75.9 51.1 90.00<br>109.3 90.00 |
| **Total reflections** | 47931 (3290) | 81322 (7264) |
| **Unique reflections** | 13200 (1003) | 11898 (1111) |
| **Multiplicity** | 3.6 (3.1) | 6.8 (6.5) |
| **Completeness (%)** | 96.5 (74.1) | 97.9 (89.2) |
| **Mean I/sigma(I)** | 11.9 (2.1) | 7.2 (1.4) |
| **R-merge** | 0.065 (0.506) | 0.171 (0.976) |
| **R-meas** | 0.089 (0.690) | 0.185 (1.062) |
| **R-pim** | 0.061 (0.467) | 0.070 (0.410) |
| **CC1/2** | 0.998 (0.718) | 0.992 (0.631) |
| **R-work** | 0.2164 (0.2984) | 0.2108 (0.3258) |
| **R-free** | 0.2421 (0.2906) | 0.2497 (0.4051) |
| **Number of non-**<br>**hydrogen atoms** | 2157 | 2026 |
| **macromolecules** | 2080 | 1991 |
| **solvent** | 77 | 30 |
| **RMS(bonds)** | 0.003 | 0.007 |
| **RMS(angles)** | 0.55 | 0.87 |
| **Ramachandran favored**<br>**(%)** | 97.08 | 97.38 |
| **Ramachandran allowed**<br>**(%)** | 2.92 | 2.62 |
| **Ramachandran outliers**<br>**(%)** | 0.00 | 0.00 |
| **Clashscore** | 2.66 | 6.07 |
| **Average B-factor** | 51.51 | 52.68 |
| **macromolecules** | 51.69 | 52.76 |
| **solvent** | 46.74 | 46.51 |
| **PDB ID** | 7SR1 | 7SR2 |

a. Statistics for the highest-resolution shell are shown in parentheses.

**Table S2.**
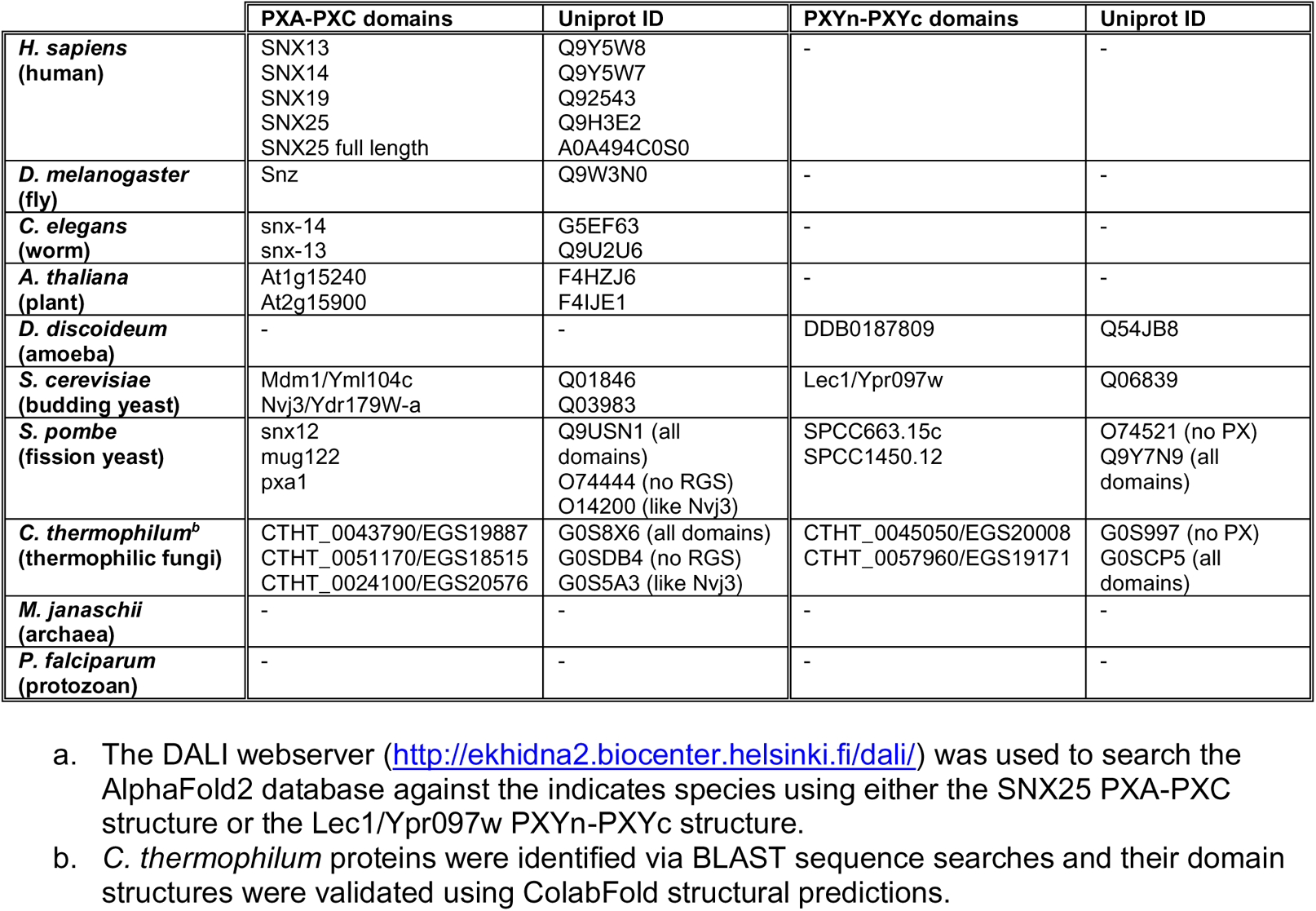
Homologues of the PXA-PXC and PXYn-PXYc domain-containing proteins based on structural similarity in the AlphaFold2 database^a^.

**Table S3.** Cavity volumes identified in LTP proteins.

| Protein | Largest continuous cavity volume (Å<sup>3</sup>)<sup>a</sup> |
| --- | --- |
| SNX13 | 2360 |
| SNX14 | 2347 |
| SNX19 | 3291 |
| SNX25 | 3974 |
| Snz | 3830 |
| Mdm1 | 2166 |
| Nvj3 | 6166 |
| Lec1/Ypr097w | 15249 |
| E-Syt2 | 1952 |
| GLTP | 380 |
| PC-TP START domain | 830 |
| CETP | 1873 |
| ORP1 | 425 |

- a. The cavity volumes in each structure were identified and calculated using POCASA (http://g6altair.sci.hokudai.ac.jp/g6/service/pocasa/)<sup>56</sup>.

**Table S4.**
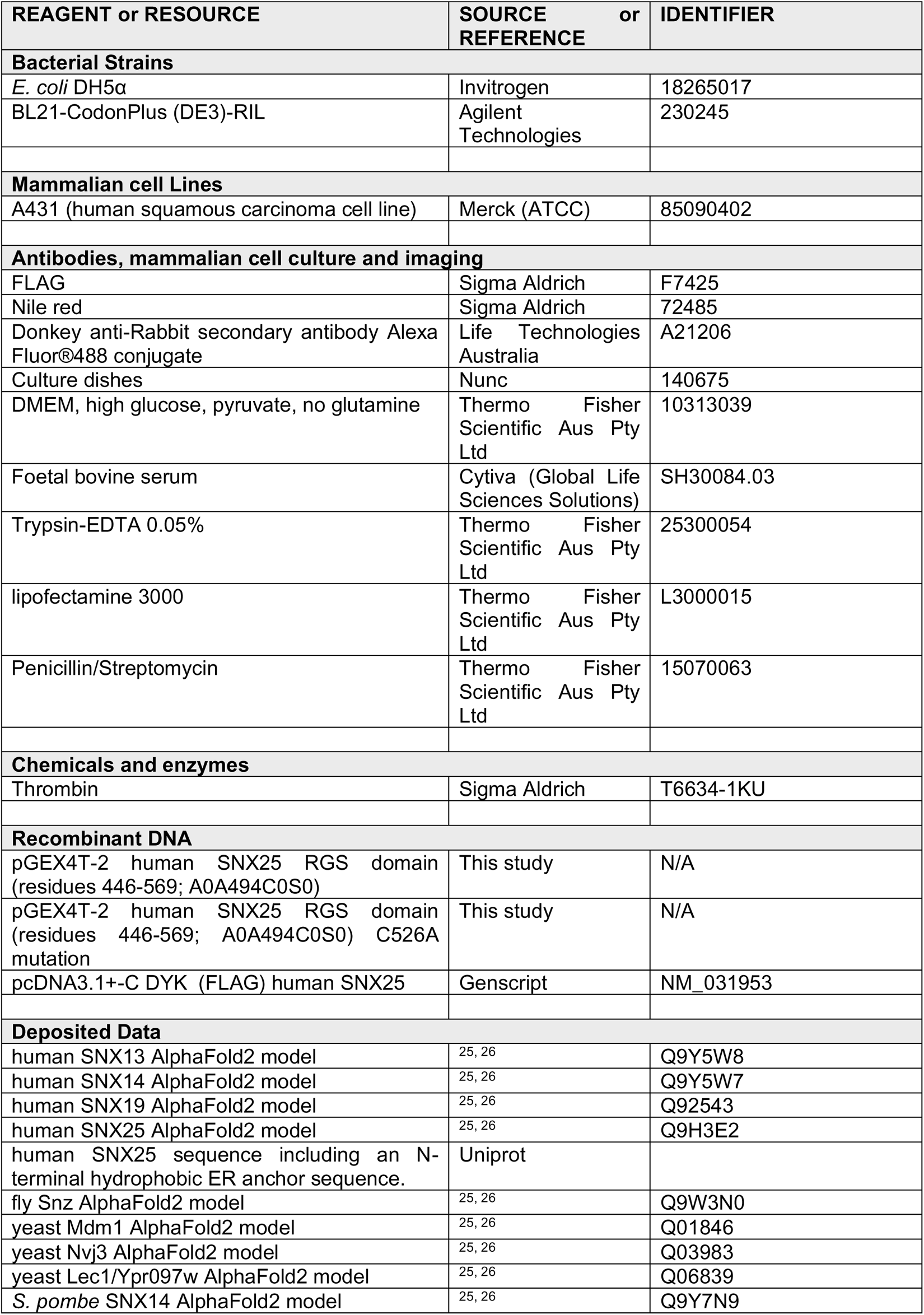
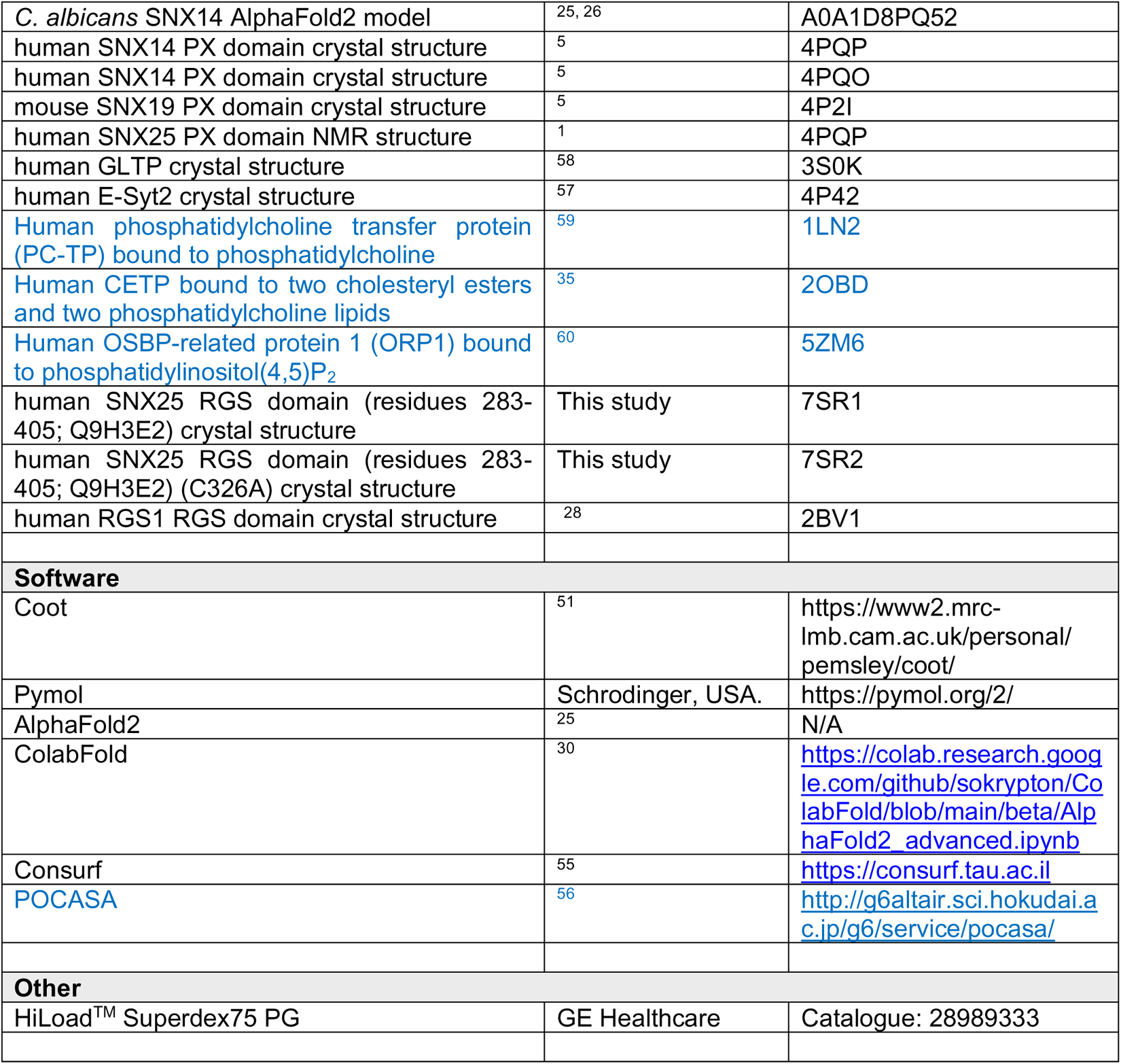
Key Resources Table.

## Notes

https://www.rcsb.org/structure/7SR1

https://www.rcsb.org/structure/7SR2

### Competing Interest Statement

The authors have declared no competing interest.

